# Medical subject heading (MeSH) annotations illuminate maize genetics and evolution

**DOI:** 10.1101/048132

**Authors:** Timothy M. Beissinger, Gota Morota

**Affiliations:** USDA-ARS Plant Genetics Research Unit, Division of Plant Sciences, MU Informatics Institute, University of Missouri, Columbia, 65211; Department of Animal Science, University of Nebraska, Lincoln, 68583

## Abstract

High-density marker panels and/or whole-genome sequencing,coupled with advanced phenotyping pipelines and sophisticated statistical methods, have dramatically increased our ability to generate lists of candidate genes or regions that are putatively associated with phenotypes or processes of interest. However, the speed with which we can validate genes, or even make reasonable biological interpretations about the principles underlying them, has not kept pace. A promising approach that runs parallel to explicitly validating individual genes is analyzing a set of genes together and assessing the biological similarities among them. This is often achieved via gene ontology (GO) analysis, a powerful tool that involves evaluating publicly available gene annotations. However, additional tools such as Medical Subject Headings (MeSH terms) can also be used to evaluate sets of genes to make biological interpretations. In this manuscript, we describe utilizing MeSH terms to make biological interpretations in maize. MeSH terms are assigned to PubMed-indexed manuscripts by the National Library of Medicine, and can be directly mapped to genes to develop gene annotations. Once mapped, these terms can be evaluated for enrichment in sets of genes or similarity between gene sets to provide biological insights. Here, we implement MeSH analyses in five maize datasets to demonstrate how MeSH can be leveraged by the maize and broader crop-genomics community.

---

Technological advances in sequencing and phenotyping have accelerated in recent decades,enabling high-throughput studies aimed at associating genotypes and phenotypes. In many cases, the speed at which we can generate large sets of candidate associations from genome-wide association studies (GWAS) (Ogura and Busch, 2015),selection mapping (Gholami et al., 2015), and other approaches has surpassed our ability to draw meaningful biological conclusions from these candidates. However, as was recently described by Rausher and Delph (2015), gene-identification is not always necessary to draw meaningful insights. Alternatively, it is often possible to look for recurrent patterns among distinct sets of candidate genes or regions in order to elucidate meaning. Annotation-based tests for enrichment or similarity represent one avenue for unraveling meaning from sets of candidates. In brief, these approaches involve identifying statistically enriched annotation terms among alist of candidate sites (usually genes or regions), or looking for similarity between terms corresponding to two sets of candidate sites, and inferring that there may be a biological explanation for the enriched or similar terms.

Commonly applied techniques often utilize gene ontology (GO) annotations (Ashburner et al., 2000), which provide putative descriptions of gene function based on automated or manual curation (Balakrishnan et al., 2013; Consortium et al., 2013). However GO term analysis isnot with out limitations. For instance, the vast majority of GO annotations are assigned algorithmically, with no human input (du Plessis et al., 2011), and the reliability of such annotations is exceeded by those that are manually curated (Škunca et al., 2012). Other areas where GO terms do not excel include that most terms are based on molecular and cellular gene products rather than macro-scale phenotypes, and due to high redundancy lists of enriched terms can be difficult to interpret (Supek et al., 2011). Therefore, despite their well-proven utility, there is growing interest in additional annotation-based approaches that can be leveraged to complement, support, enhance, or add to the patterns identified by GO. Included among this assortment of strategies are KEGG annotations (Kanehisa and Goto, 2000), Disease Ontology (Schriml et al., 2012), and Medical Subject Headings (MeSH), which were introduced at the National Library of Medicine (NLM) more than fifty years ago (Lipscomb, 2000).

MeSH terms are the NLM’s controlled terminology, primarily used to organize and index information and manuscripts found in common databases such as PubMed https://www.nlm.nih.gov/mesh/meshhome.html.By mapping from MeSH terms to manuscripts, and then to a list of candidate genes, a semantic patternsearch for biological meaning can be conducted (Nakazato et al., 2008). Recently, the MeSH Over-representation Analysis (ORA) Framework, a suite of software for conducting MeSH enrichment analyses using R (R Core Team, 2015) and Bioconductor (Huber et al., 2015), was developed (Tsuyuzaki et al., 2015). MeSH analysis has proven useful for deducing meaning from sets of genes implicated across several agricultural animal species including in cattle, swine, horse and chicken(Morota et al., 2016, Morota et al., 2015). Here, we implement five MeSH analyses in maize, which collectivelydemonstrate how MeSH can been used to enrich biological understanding in crop species.

In this study, which is meant to be both a primer for MeSH-based analysis in maize and other crop plants, as well as an investigation of patterns that can be deduced regarding maizegenetics and evolution, we identify over-represented MeSH terms among candidate genes identified from five distinct maize datasets: 1) regions under selection during maize domestication (Hufford et al., 2012); 2) regions under selection during maize improvement (Hufford et al., 2012); 3) regions under selection for seed size (Hirsch et al., 2014); 4) regions under selection for ear number (Beissinger et al., 2014); and 5) regions contributing to inflorescence traits (Brown et al., 2011). After identifying significant MeSH terms, we also assess and test for semantic similarity, or MeSH-based relatedness, among the genes identified in each of these datasets to identify relationships among the genetic underpinnings of these traits/selection regimes.

## Materials and methods

### Code availability

To enable implementation of MeSH analyses by other researchers, all scripts used in this study are available as annotated supplemental files in R-markdown format (Supplemental files S1 - S7). Scripts were written inR (R Core Team, 2015) and utilize Bioconductor (Huber et al.,2015), the MeSH ORA Framework including the “meshr” for ORA and the “mesh.zma.eg.db” maize-specific mapping table (Tsuyuzaki et al., 2015), and MeSHSim (Zhou and Shui, 2015). The mapping table provides the necessary link between NCBI Entrez IDs and NLM MeSH IDs. For maize, the mapping table was generated using both gene2pubmed ftp://ftp.ncbi.nih.gov/gene/DATA/. with data licensed by PubMed and a reciprocal BLAST best hit search conducted across 115 organisms and requiring an E-value of 50, with additional details described by Tsuyuzaki et al (2015). The GOstats R package (Falcon and Gentleman, 2007) was used to implement GO ORA to generate a baseline that MeSH results could be compared to. Genome data was downloaded using the biomaRt R package (Durinck et al, 2009). Full analysis details are included within the reproducible scripts (Supplemental files S1-S7), so here we provide an overview of the data and analyses implemented.

### Datasets

We analyzed five publicly available datasets to identify enriched MeSH terms and look for semantic similarity between different traits and selection regimes. The datasets analyzed are described in Table 1. For the four datasets that involved contiguous regions (Domestication, improvement, seed size,and ear number), all genes that fell within the implicated regions were used for MeSH analysis. For the remaining dataset (inflorescence traits), which involved isolated SNPs identified through GWAS instead of genomic regions, all genes within 10kb of the implicated SNPs were used for MeSH analysis. All genemodels and gene locations were based on the maize reference genome version 2 (Schnable et al., 2009).

**Table 1.** This table describes the datasets used in this study, including reference information where full details can be found and a brief description of each.

| **Dataset** | **Reference** | **Description** |
| --- | --- | --- |
| **Domestication** | Hufford et al, 2012 | Regions selected during domestication from teosinte to maize. |
| **Improvement** | Hufford et al, 2012 | Regions selected during post-domestication maize improvement. |
| **Seed Size** | Hirsch et al, 2014 | Regions artificially selected for seed size in a long-term selection experiment. |
| **Ear Number** | Beissinger et al, 2014 | Regions artificially selected for ear number in a long-term selection experiment. |
| **Inflorescence Traits** | Brown et al, 2011 | SNPs associated with inflorescence traits from a genome-wide association study. |

### Analyses

Each of the five datasets was first tested for any over-represented MeSH terms and GO terms. MeSH ORA was performed using the MeSH ORA Framework which includes the “meshr” and “MeSH.Zma.eg.db” R-packages (Tsuyuzaki et al., 2015), the latter of which is a mapping table that connects gene Entrez IDs to MeSH IDs. These packages can be installed using Bioconductor by running the command, “source(“https://bioconductor.org/biocLite.R”)”, followed by “biocLite(“meshr“)” and “biocLite(“MeSH.Zma.eg.db“)”. Further instructions to install and run these packages are provided in supplemental files S1 - S5. Unfortunately, the majority of maize genes annotated in the maize version 2 reference genome (Schnable et al, 2009) do not have a corresponding Entrez ID, and therefore are not useful for MeSH analyses. Of the 40,481genemodelsavailable from Ensembl Plants (http://plants.ensembl.org/index.html), only14,142 havecorresponding Entrez IDs. The “meshHyperGTest” function was implemented to conduct a hypergeometric test. Specifically, to test the probability that a specific MeSH term is enriched in a particular set of genes, as compared to a background gene set, this function calculates where N is the total number of background genes, k is the number of genes in the set being tested, M is the number of background genes corresponding to the particular MeSH term, and s is the number of genes in the testset that correspond to that MeSH term (Tsuyuzaki et al., 2015). For this study, all Entrez genes inthe maize reference genomeversion 2 (Schnable et al., 2009) were used as the background gene set. GO ORA was conducted using a similar approach, as demonstrated in the supplemental files. The necessary GOstats package is installed by running “biocLite(“GOstats”)”.

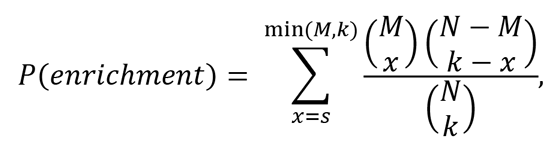

Next, semantic similarity between distinct experiments was evaluated using the MeSHSim R package (Zhou and Shui, 2015) to elucidate if there are underlying relationships between the trait data-sets (seed size, ear number, or inflorescence traits) and the process datasets (domestication, improvement), as well as the relationships within the process and trait datasets. The “headingSetSim” function was used, and results were plotted with the corrplot R package (Wei, 2013).

## Results

### Overrepresentation analysis

MeSH ORA involves performing a hypergeometric test to determine which MeSH terms are enriched among the candidate set of genes compared to a set of background genes. Allgenes in the maize reference genome version 2 (Schnable et al., 2009) with Entrez IDs were used as the background set. While GO terms are classified into the three groups “molecular function”, “cellular components”, and “biological processes”, MeSH classifications include several groups, many of which are geared more toward indexing biomedical manuscripts than biological processes. However, classifications including “chemicals and drugs”, “diseases”, “anatomy”, and “phenomena and processes”, all have the potential to contribute to the biological understanding of sets of genes. Overrepresented terms in each of these categories for the five analyzed datasets are described in Supplemental Files S1 - S5. For the purpose of demonstration, MeSH terms identified within the “anatomy” classification are provided as an example and described in detail in Table 2. Many of the enriched terms serve to provide additional evidence for reasonable a priori expectations, such as the observation that “flowers” and “seeds” are both enriched within the set of genes under selection during domestication. However, others introduce interesting questions that could serve to drive hypothesis generation for future studies. For instance, the only enriched term identified from the ear number dataset is “endosperm”, which one would not immediately assume to be related to ear number.

**Table 2.**
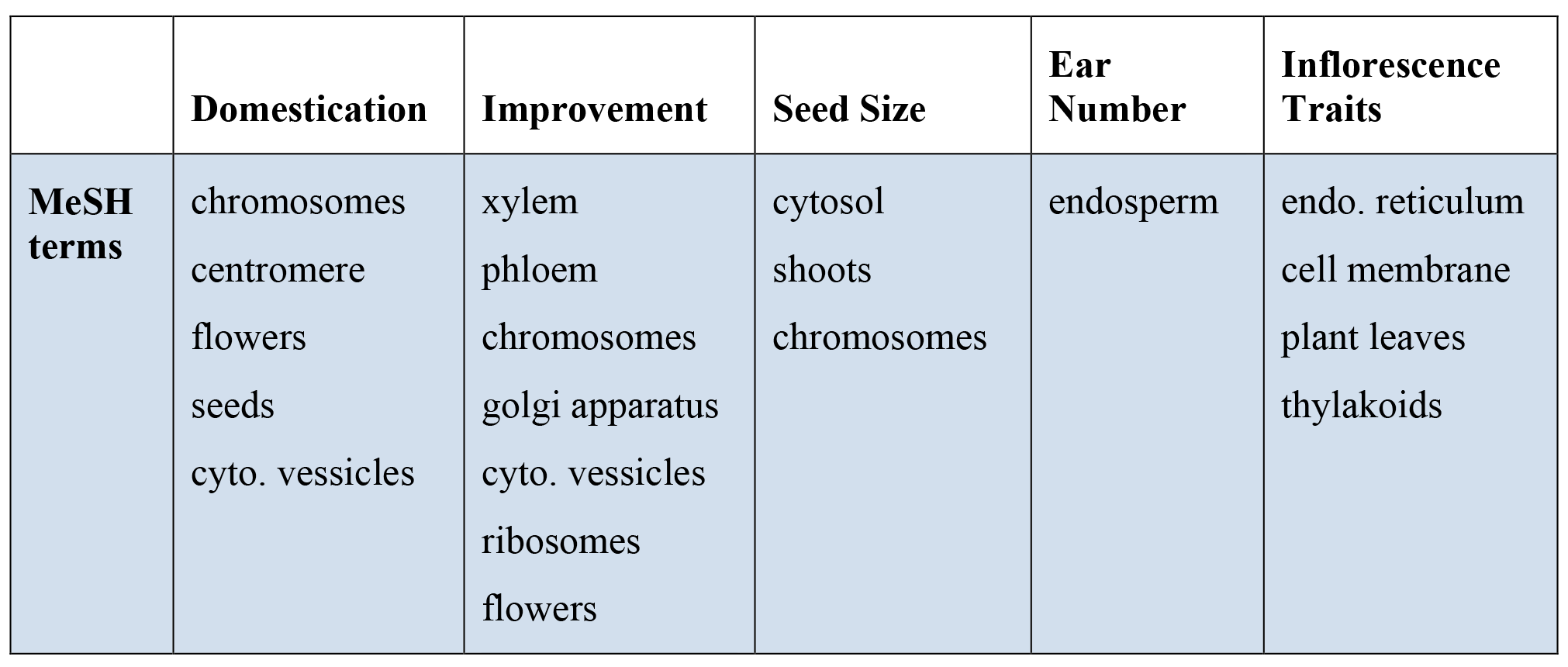
MeSH terms enriched in each of the five datasets within the anatomy MeSHclassification group.

|  | **Domestication** | **Improvement** | **Seed Size** | **Ear Number** | **Inflorescence Traits** |
| --- | --- | --- | --- | --- | --- |
| **MeSH terms** | chromosomes<br>centromere<br>flowers<br>seeds<br>cyto. vessicles | xylem<br>phloem<br>chromosomes<br>golgi apparatus<br>cyto. vessicles<br>ribosomes<br>flowers | cytosol<br>shoots<br>chromosomes | endosperm | endo. reticulum<br>cell membrane<br>plant leaves<br>thylakoids |

### Semantic similarity analysis

Another powerful use of MeSH is that it can be used to calculate the semantic similarity between distinct sets of MeSH terms. This type of analysis enables one to look for hidden relationships among sets of genes, potentially uncovering biological meaning. For the five datasets we studied, we assessed whether there were pairwise relationships linking any of them. Figure 1 depicts the MeSH similarity between each set of candidate genes. Interestingly, the strongest relationship identified was between domestication genes and seed size genes, possibly suggesting that seed size traits were more strongly selected during domestication than were ear number or other inflorescence traits. Noteworthy relationships were also observed between domestication and improvement genes, as well as between seed size and improvement genes. It should be noted that ear number genes were not strongly related to any ofthe other gene sets, which may simply result from the fact that the ear number dataset included the fewest candidate genes. This possibility is elaborated upon further inthediscussion.

**Figure 1.**
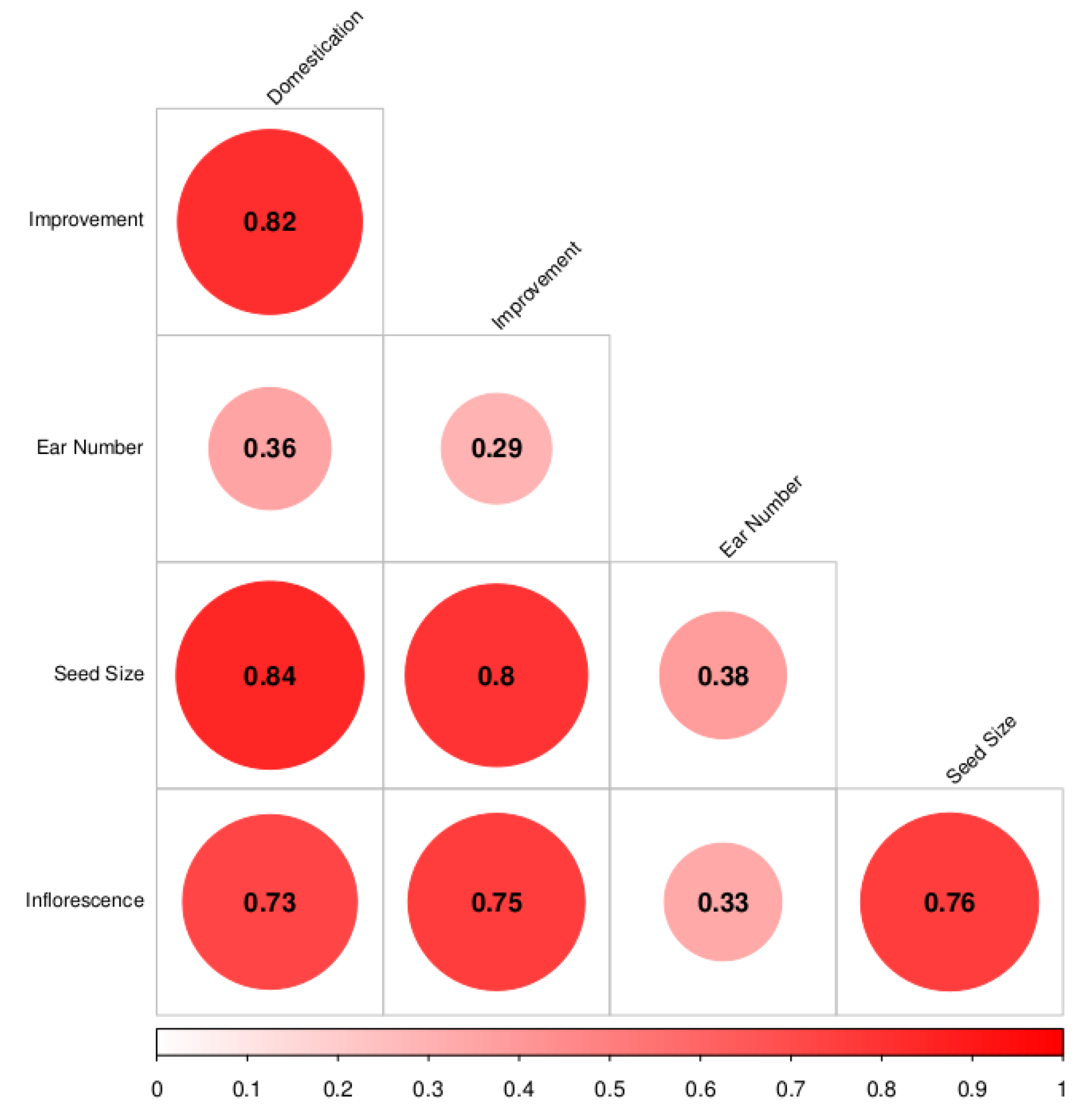
MeSH semantic similarity-based relatedness among sets of genes implicated in each of the five datasets studied. The size of each circle, degree of red shading, and value reported correspond to the relatedness between each pair of datasets.

### Comparison of real data to a random set of genes

We conducted an analysis of 1,500 randomly selected genes to determine the robustness of MeSH analyses in a scenario where no biological meaning is present (SupplementalFile S6). As is expected for any p-value based method, a subset of terms achieved significance. Spurious results were also observed in a parallel GO analysis (Supplemental File S6). In contrast to many of the real datasets we evaluated, there was no overwhelming theme tying the terms together. This subjective observation is supported by a semantic similarity analysis between the random gene set and the real datasets, where lower similarities were generally observed (Supplemental File S7). Still, the observation that “significant” MeSH or GO terms can arise from a random set of genes suggests that caution should be exercised when attempting to make interpretations from any such study, as is discussed in detail by Pavlidis et al. (2012). Although we utilized a lenient p=0.05 significance threshold here, in part for the purpose of demonstration, the use of a hypergeometric distribution for testing allows a more stringent significance threshold to be employed when needed.

## Discussion

Our analysis of five existing datasets demonstrates how MeSH ORA and semantic-similarity analyses can be used to mine data and confirm and/or generate informative hypotheses. Like GO, MeSH-based approaches leverage curated annotations to provide biological insights. However, in contrast to GO, MeSH curations are manually assigned by the NLM, and therefore they have the potential to be more accurate and interpretable. In fact, as we have shown, several of the enriched terms within the #x201C;anatomy#x201D; category are directly related to macrophenotypes, such as “seeds”, “shoots”, “flowers”, and “ears”. Whether applied to existing data, as we have demonstrated here, or if used to infer meaning from a list of candidates generated from a novel mapping study, MeSHrepresents an additional toolfor drawing inferences from large-scale sets of genomic data.

### Biological implications

Among the findings gleaned from this analysis was the observation that while both “flowers” and “seeds” were enriched terms in the domestication set of genes, only “flowers” remained significant among improvement genes (Table 2). This result is consistent with the morphological observation that the maize female inflorescence is dramatically different from that of teosinte (Gottlieb, 1984), with one of the most immediately apparent differences being seed related; the teosinte outer glume forms a hard teosinte fruitcase that completely encapsulates each kernel, while in maize the outer glume is barely present (Dorweiler and Doebley, 1997). It has been shown that this trait is controlled by relatively few genes, with tga1 (Wang et al., 2005, Wang et al., 2015) being of particular importance, and therefore our MeSH finding may suggest that after intense selection on seed traits during domestication, subsequent selection on further seed modifications during improvement has possibly been more subdued.

The hypothesis that domestication immediately impacted seed-related traits more than others is further supported by our semantic similarity analysis, where the most similar pair of gene-sets we tested corresponded to domestication and seed size (Figure 1). Also, while the limited number of genes included in the ear-number dataset (Beissinger et al., 2014) seems to constrain the estimated similarity between ear-number genes and the other datasets, we do observe that ear-number genes are semantically more similar to domestication genes than they are to improvement genes (Figure 1). This again is consistent with morphological differences between maize and teosinte, with maize demonstrating apical dominance while teosinte has a much more branched structure (Doebley et al.,1997). The observation of greater similarity between ear number genes and domestication genes than between ear number genes and improvement genes lends support to the existing supposition that single-eared plants have likely been favorable throughout the era of post-domestication maize improvement due to the ease with which such plants can be hand harvested (De Leon and Coors, 2002).

An observation that ran contrary to our expectation was that “shoots”was an enriched term among seed size genes, while “endosperm” was enriched within the set of ear number genes (Table 2). We are tempted to dismiss these findings as spurious, but both have plausible biological explanations. In the Krug selection population (Hirsch et al., 2014), where our seed size regions were identified, mass selection not only impacted seed size, but also affected seedling size, leaf width, stalk circumference, and cob weight (Sekhon et al., 2014), indicating that the set of genes selected for seed size also being implicated in shoot traits is not unexpected. Similarly, the ear number genes were identified from the Golden Glow selection experiment for ear number (Maita and Coors, 1996), where correlated changes in kernel size and kernel number were also observed (De Leon and Coors, 2002).

### Comparison of MeSH and GO Overrepresentation Analyses

Among the most obvious findings when comparing results from MeSH and GO for all five of the datasets is that the number of GO term associations dramatically surpasses the number identified by MeSH. This can be observed in Supplemental Files S1-S5, which contain lists of associations from the implementation of both methods. Within these lists, there are cases of clearly overlapping GO and MeSH terms. For instance, in the improvement dataset, MeSH identified “Lipoxygenase” as the most significantly overrepresented term in the Chemicals and Drugs category, while GO identified the similar “linoleate 13S-lipoxygenase activity” term as highly significant in the Molecular Function category. However, there were instances where the literature-based nature of MeSH allowed it to identify associations that were easily missed by GO.An example of this is that from the inflorescence dataset “Hybrid Vigor” was an enriched term in the Phenomena and Processes MeSH category, while no similar terms were identified by GO in any category. Although these examples are anecdotal, they are only a minor subset of the complete lists provided by this analysis and available for further scrutiny (Supplemental Files S1-S5). We mention the examples to demonstrate that MeSH and GO can either differ remarkably in their findings or in some instances, particularly for highly significant terms. provide an independent confirmation that the other method is on the right track.

### Current Limitations

Despite the promising MeSH ORA and semantic similarity results observed in this study, using MeSH to guide biological interpretations still has an assortment of limitations that should be considered during any study that involves MeSH. Firstly, for non-model organisms, including maize and other crops, relatively few genes have corresponding manuscripts that have been directly annotated with MeSH terms. Instead, the MeSH.Zma.eg.dbR package/mapping table also relies on mapping genes to model species based on reciprocal BLAST best hits, and gleaning MeSH terms from there (Tsuyuzaki et al., 2015). Additionally, a requirement of current software isthat all genes have Entrez gene ID’s (Maglott et al., 2005) to enable mapping from genes to MeSH terms, but Entrez ID’s have only been assigned to a subset of maize genes. In fact, among the five datasets we analyzed, approximately two thirds of the genes falling within the putatively functional regions did not have a corresponding Entrez ID. This is particularly troubling in light of our observations regarding the ear number gene set, which was the smallest list of genes considered. Only 195 genes were contained within the selected regions (compared to thousands for some of the other data sets), and only 62 of those had corresponding Entrez IDs. With fewer genes included during ORA, the power to detect significant enrichment is reduced. Similarly, this dataset showed very weak similarity to the others, which we hypothesize is at least in part due to the limited number of included genes and corresponding MeSH terms.

Even considering these limitations, we expect MeSH-based analyses will improve overtime. As additional mapping and functional manuscripts are published, the number of Entrez genes and the descriptive MeSH terms corresponding to each, in both model and non-model species, will increase. This increase will improve the magnitude and reliability of results gleaned from MeSH. Although improvements are expected with time, the five datasets studied here demonstrate how MeSH can currently be leveraged for making biological interpretations in maize as well as other crop species.

## Acknowledgments

This work was supported by the USDA Agricultural Research Service. A University of Nebraska Layman fund provided support for Gota Morota.

## References

M., Ashburner, C.A. Ball, J.A. Blake, D. Botstein, H. Butler, J.M. Cherry, A.P. Davis, K. Dolinski,S.S. Dwight, J.T. Eppig, et al., 2000. Gene ontology: tool for the unification of biology. Nature genetics 25:25–29.

R., Balakrishnan, M.A. Harris, R. Huntley, K. Van Auken, and J.M. Cherry, 2013. A guide to best practices for gene ontology (go) manual annotation. Database 2013:bat054.

T.M., Beissinger, C.N. Hirsch, B. Vaillancourt, S. Deshpande, K. Barry, C.R. Buell, S.M. Kaeppler, D. Gianola, and N. de Leon, 2014. A genome-wide scan for evidence of selection in a maize population under long-term artificial selection for ear number. Genetics 196:829–840.

P.J., Brown, N. Upadyayula, G.S. Mahone, F. Tian, P.J. Bradbury, S. Myles, J.B. Holland, S. Flint-Garcia, M.D. McMullen, E.S. Buckler, et al., 2011. Distinct genetic architectures for male and female inflorescence traits of maize. PLoS Genet 7:e1002383.

G.O. Consortium et al., 2013.Gene ontology annotations and resources. Nucleic acids research 41:D530–D535.

De Leon N., and J. Coors, 2002. Twenty-four cycles of mass selection for prolificacy in the golden glow maize population. Crop science 42:325–333.

Doebley, J., A. Stec and L. Hubbard, 1997. The evolution of apical dominance in maize. Nature 386:485–488.

Durinck, S., Spellman P.T., Birney E.,, and Huber, W., 2009. Mapping identifiers for the integration of genomic datasets with the R/Bioconductor package biomaRt. Nature Protocols 4:1184–1191.

Dorweiler, J. and J. Doebley, 1997. Developmental analysis of teosinte glume architecture!: A key locus in the evolution of maize (poaceae). American Journal of Botany 84:1313–1313.

Falcon, S., and Gentleman, R., 2007. Using GOstats to test gene lists for GO term association. Bioinformatics 23:257–8.

Gholami, M., C. Reimer, M. Erbe, R. Preisinger, A. Weigend, S. Weigend, B. Servin and H. Simianer 2015. Genome scan for selection in structured layer chicken populations exploiting linkage disequilibrium information. PloS one 10:e0130497.

Gottlieb, L., 1984. Genetics and morphological evolution in plants. AmericanNaturalist.Pp681–709.

Hirsch, C.N., S.A. Flint-Garcia, T.M. Beissinger, S.R. Eichten, S. Deshpande, K. Barry, M.D. McMullen, J.B. Holland, E.S. Buckler, N. Springer, et al., 2014. Insights into the effects of longterm artificial selection on seed size in maize. Genetics 198:409–421.

Huber, W., Carey, V.J., Gentleman, R., Anders, S., Carlson,M., Carvalho, B.S., Bravo, H.C., Gatto, L., Girke, T., Gottardo, R., Hahne, F., Hansen, K.D., Irizarry, R.A., Lawrence, M., Love, M.I., MacDonald, J., Obenchain, V., Ole’s, A. K., Pag‘es, H., Reyes, A., Shannon, P., Smyth,G.K., Tenenbaum, D., Waldron, L., Morgan, and M., 2015. Orchestrating high-throughput genomic analysis with Bioconductor. Nature Methods 12:115–121. http://www.nature.com/nmeth/journal/v12/n2/full/nmeth.3252.html.

Hufford, M.B., X. Xu, J. Van Heerwaarden, T. Pyhäjärvi, J.-M. Chia, R.A. Cartwright, R.J. Elshire, J.C. Glaubitz, K.E. Guill, S.M. Kaeppler, et al., 2012. Comparative population genomics of maize domestication and improvement. Nature genetics 44:808–811.

Kanehisa, M. and S. Goto, 2000. Kegg: kyoto encyclopedia of genes and genomes. Nucleic acids research 28:27–30.

Lipscomb, C.E., 2000. Medical subject headings (MeSH). Bulletin of the Medical Library Association 88:265.

Maglott, D., J. Ostell, K.D. Pruitt, and T. Tatusova, 2005. Entrez gene: gene-centered information at ncbi. Nucleic acids research 33:D54–D58.

Maita, R. and J. Coors, 1996. Twenty cycles of biparental mass selection for prolificacy in the open-pollinated maize population golden glow. Crop science 36:1527–1532.

Morota, G., T.M. Beissinger, and F. Peñagaricano, 2016. MeSH annotation of the chicken genome: Mesh-informed enrichment analysis and MeSH-guided semantic similarity among functional terms and gene products. bioRxiv P.034975.

Morota, G., F. Peñagaricano, J.L. Petersen, D.C. Ciobanu, K. Tsuyuzaki, and I. Nikaido, 2015. An application of MeSHF enrichment analysis in livestock. Animal Genetics 46:381–387. http://dx.doi.org/10.1111/age.12307

Nakazato, T., T. Takinaka, H. Mizuguchi, H. Matsuda, H. Bono, and M. Asogawa, 2008. Biocompass: a novel functional inference tool that utilizes MeSH hierarchy to analyze groups of genes. In silico biology 8:53–61.

Ogura, T. and W. Busch, 2015. From phenotypes to causal sequences: using genome wide association studies to dissect the sequence basis for variation of plant development. Current opinion in plant biology 23:98–108.

Pavlidis, P., J.D. Jensen, W. Stephan, and A. Stamatakis, 2012. A critical assessment of storytelling: gene ontology categories and the importance of validating genomic scans. Molecular biology and evolution 29:3237–3248.

du Plessis, L., N. Škunca, and C. Dessimoz, 2011. The what, where, how and why of gene ontologya primer for bioinformaticians. Briefings in bioinformatics P.bbr002.

R Core Team, 2015. R:A Language and Environment for Statistical Computing. R Foundation for Statistical Computing, Vienna,Austria. https://www.R-project.org/.

Rausher, M.D. and L.F. Delph, 2015. Commentary:When does understanding phenotypic evolution require identification of the underlying genes? Evolution 69:1655–1664.

Schnable, P.S., D. Ware, R.S. Fulton, J.C. Stein, F. Wei, S. Pasternak, C. Liang, J. Zhang, L. Fulton, T.A. Graves, et al., 2009. The b73 maize genome:complexity, diversity, and dynamics. science 326:1112–1115.

Schriml, L. M., C. Arze, S. Nadendla, Y.-W.W. Chang, M. Mazaitis, V. Felix, G. Feng, and W.A. Kibbe, 2012. Disease ontology: a backbone for disease semantic integration. Nucleic acids research 40:D940–D946.

Sekhon, R.S., C.N. Hirsch, K.L. Childs, M.W. Breitzman, P. Kell, S. Duvick, E.P. Spalding, C.R. Buell, N. de Leon, and S.M. Kaeppler, 2014.Phenotypic and transcriptional analysis of divergently selected maize populations reveals the role of developmental timing in seed size determination. Plant physiology 165:658–669.

Supek, F., M. Bošnjak, N. Škunca, and T. Šmuc, 2011. Revigo summarizes and visualizes long lists of gene ontology terms. PloS one 6:e21800.

Tsuyuzaki, K., G. Morota, M. Ishii, T. Nakazato, S. Miyazaki, and I. Nikaido, 2015. Mesh ora framework: R/bioconductor packages to support mesh over-representation analysis. BMC bioinformatics 16:45.

Škunca, N., A. Altenhoff, and C. Dessimoz, 2012. Quality of computationally inferred gene ontology annotations. PLoS Comput Biol 8:1–11. http://dx.doi.org/10.1371%2Fjournal.pcbi.1002533

Wang, H., T. Nussbaum-Wagler, B. Li, Q. Zhao Y. Vigouroux, M. Faller, K. Bomblies, L. Lukens, and J.F. Doebley, 2005. The origin of the naked grains of maize. Nature 436:714–719.

Wang, H., A.J. Studer, Q. Zhao, R. Meeley, and J.F. Doebley, 2015. Evidence that the origin of naked kernels during maize domestication was caused by a single amino acid substitution in tga1. Genetics 200:965–974.

Wei, T., 2013. corrplot: Visualization of a correlation matrix. https://CRAN.R-project.org/package=corrplot. R package version 0.73.

Zhou, J. and Y. Shui, 2015. MeSHSim: MeSH(Medical Subject Headings) Semantic Similarity Measures. R package version 1.2.0.

